# Divide and Cluster: The DIVINE Framework for Deterministic Top-Down Analysis of Molecular Dynamics Trajectories

**DOI:** 10.1101/2025.06.20.660828

**Authors:** Jherome Brylle Woody Santos, Lexin Chen, Ramón Alain Miranda-Quintana

## Abstract

We present DIVIsive *N*-ary Ensembles (DIVINE), a deterministic, top-down clustering framework designed for molecular dynamics (MD) trajectories. DIVINE constructs a complete clustering hierarchy by recursively splitting clusters based on *n*-ary similarity principles, avoiding the need for O(*N*^2^) pairwise distance matrices. It supports multiple cluster selection criteria, including a weighted variance metric, and deterministic anchor initialization strategies such as NANI (*N*-ary Natural Initiation), ensuring reproducible and structurally meaningful partitions. Testing DIVINE up to a 305 μs folding trajectory of the villin headpiece (HP35) revealed that it matched or exceeded the clustering quality of bisecting *k*-means while reducing runtime and eliminating stochastic variability. Its single-pass design enables efficient exploration of clustering resolutions without repeated executions. By combining scalability, interpretability, and determinism, DIVINE offers a robust and practical alternative to conventional MD clustering methods. DIVINE is publicly available as part of the MDANCE package: https://github.com/mqcomplab/MDANCE.

## INTRODUCTION

Molecular dynamics (MD) simulations have become an important tool in the exploration of structure and dynamics of biomolecular systems at atomic resolution.^1,2^ With the rise of GPU acceleration and broader access to high-performance computing, it is now routine for simulations to reach micro-to millisecond timescales.^3–5^ These simulations generate massive, high-dimensional datasets comprising millions of conformational snapshots, which require careful and robust post-processing techniques to extract chemically meaningful information. A central task in this process is clustering, which involves the identification of discrete structural states, estimation of their populations, and visualization of their structural relationships.^6–10^

Traditional clustering pipelines used in MD analysis generally fall into two main categories: partitional methods such as *k*-means, and hierarchical methods, such as hierarchical agglomerative clustering (HAC). *k*-means is widely popular due to its computational efficiency and scalability. However, standard *k*-means assumes convex shapes, essentially biasing partitions toward globular clusters separated by linear decision boundaries. This assumption could be problematic for high-dimensional MD trajectories, where conformational states often form complex, curved, or elongated shapes that cannot be correctly separated by such simple geometric rules. Additionally, *k*-means can be quite sensitive to initialization, which limits its reliability and reproducibility when applied to MD data.^11,12^ To address some of these issues, we previously developed *N*-ary Natural Initiation (NANI), a deterministic initialization scheme that improves clustering compactness and stability.^13^

Beyond initialization, however, the sheer computational cost of standard *k*-means on massive datasets could present another challenge. To address this bottleneck, Sculley proposed the Mini-Batch *k*-means algorithm.^14^ By utilizing small, random batches of data to iteratively update cluster centroids, this method drastically reduces memory usage and convergence time, which effectively enables large-scale clustering.^15,16^ Yet, despite the efficiency gains, Mini-Batch *k*-means retains the fundamental topological limitations of the *k*-means algorithm: it still produces a flat partition that forces data into convex-shaped clusters. Moreover, the stochastic nature of the mini-batch updates introduces additional variance into the optimization process, which can interact with initialization effects and lead to variability in the results. As a consequence, multiple runs are often employed in practice to assess stability, highlighting an inherent trade-off between computational efficiency and reproducibility in large-scale *k*-means-based clustering.

HAC, on the other hand, constructs a dendrogram by iteratively merging the most similar frames, allowing the identification of arbitrarily shaped clusters. However, its O(*N*^2^) time and memory complexity from the need to compute and store all pairwise distances make it impractical for large MD datasets.^7,17^ To address this, we proposed Hierarchical Extended Linkage Method (HELM), a hybrid clustering protocol that uses *k*-means NANI for efficient pre-clustering, followed by agglomerative merging.^18^ While this approach retained much of HAC’s flexibility without a full distance matrix, it still relies on an agglomerative (bottom-up) strategy.

As an alternative, divisive clustering offers a top-down framework. It starts with the entire dataset as a single cluster, which is recursively split.^19^ This strategy naturally captures global structure and crucially avoids ever constructing a full pairwise distance matrix if splits and decisions are made based on cluster-wise properties. A notable example is the bisecting *k*-means^20,21^ (referred to as BKM in this paper), which iteratively splits the cluster with the biggest inertia (sum of squared errors) or largest number of frames using a standard *k*-means (with *k* = 2). While efficient, common implementations produce only a flat partition unless explicitly modified to retain intermediate splits. Moreover, the reliance on random or *k*-means++ initialization compromises reproducibility and requires multiple runs to properly assess clustering stability.

Another example is DIvisive ANAlysis (DIANA)^22^, a classical top-down method. DIANA selects the cluster with the largest diameter (i.e., the most internally dissimilar cluster) and splits it by isolating a “splinter” group.^23^ While conceptually appealing, DIANA also requires an O(*N*^2^) dissimilarity matrix making it not feasible for large-scale MD data. To our knowledge, no MD analysis-specific library currently implements a fully divisive clustering approach for millions of conformational frames.

In this work, we present DIVIsive *N*-ary Ensembles (DIVINE), a deterministic top-down clustering framework designed for large-scale MD simulations. DIVINE begins with all frames in a single cluster and recursively splits them using *n*-ary similarity principles, enabling efficient O(*N*) cluster evaluations without requiring pairwise comparisons.^13,18,24–34^ We incorporate three criteria for selecting clusters to split (mean squared deviation (MSD), radius, and weighted MSD) and three anchor-selection strategies (NANI-based, outlier-based, and DIANA-like splintering). Every step in DIVINE is fully deterministic, ensuring that identical inputs produce exactly the same outputs.

We benchmark DIVINE on a folding trajectory of the villin headpiece (HP35), comparing its performance against scikit-learn’s BKM.^20^ Our results demonstrate that DIVINE produces clear hierarchical clustering that directly maps conformational lineages while enabling efficient assessment of clustering quality in a single pass. When evaluated using standard validation metrics like Calinski–Harabasz^35^ and Davies– Bouldin^36^ indices, DIVINE matches or outperforms BKM, with superior runtime stability and full reproducibility. Unlike conventional divisive methods, DIVINE scales efficiently while preserving hierarchy, allowing rapid exploration of clustering resolutions and making it a powerful addition to MD analysis pipelines. DIVINE is publicly available as part of the MDANCE package: https://github.com/mqcomplab/MDANCE.

## THEORY AND IMPLEMENTATION

### Clustering Framework Overview

DIVINE begins by treating the entire trajectory (consisting of N frames) as the initial cluster. It then proceeds iteratively: at each step, one cluster is selected for division and is then split into two subclusters. The process continues until a termination criterion is met.

Two stopping modes are supported: (a) end = “points”, which continues until every cluster contains a single frame (i.e., an exhaustive hierarchy down to singletons), and (b) end = “*k*”, which stops when a specified number of clusters *k* is reached. Users may choose end = “*k*” for a predetermined clustering or end = ‘points’ to extract the full dendrogram and then decide an optimal *k* by analyzing clustering metrics.

To suppress formation of unwanted, low-population clusters, we introduce an optional threshold parameter that enforces a minimum size for valid subclusters. If a proposed split results in any subcluster with fewer than threshold × N frames, that split is skipped. This prevents overfragmentation as we treat extremely rare outlier clusters as noise.

### Cluster Selection Criteria

At each iteration, DIVINE must decide which cluster to split. We provide three selection criteria via the split parameter. We also note that while we describe these metrics below in the context of Cartesian coordinates, the framework is feature-agnostic: these criteria apply equally to alternative input features such as dihedral angles, pairwise distances, and other collective variables.

#### MSD (Mean Squared Deviation)

Selects the cluster with the greatest internal dispersion as measured by the MSD of its frames. The MSD is the average of the pairwise squared distances computed in linear time (O(*N*)) using a cluster summary representation. A higher MSD indicates a structurally diverse cluster and thus a strong candidate for further subdivision. This criterion is similar in spirit to DIANA’s diameter but uses all points collectively rather than just the single furthest pair, making it less sensitive to an isolated outlier.

#### radius

Selects the cluster with the largest radius, defined as the maximum distance between the cluster’s medoid (most representative frame) and any other frame within the cluster. The medoid is determined using complementary MSD (cMSD), a measure of the frame’s contribution to the overall cluster dispersion. For each frame in the cluster, cMSD is computed as the MSD of all other frames when that frame is excluded. The frame whose exclusion results in the largest increase in dispersion (i.e., the highest cMSD) is considered the most central to the cluster and is selected as the medoid. The radius is then defined as the largest distance between this medoid and any other frame in the cluster.

#### weighted_MSD

Choose the cluster maximizing MSD × m (where m is the cluster’s size). This weighted variance criterion biases the selection toward large clusters that are also dispersed. By incorporating cluster size into the decision metric, this directs the focus toward larger clusters that account for a greater share of the system’s total structural variation.

### Anchor Frame Selection

Once a cluster is selected for splitting, DIVINE now chooses two anchor frames to seed the resulting subclusters. We implemented multiple anchor selection strategies:

#### NANI

This mode leverages the *k*-means NANI algorithm to identify two high-quality initial centers for splitting. NANI selects diverse frames from high-density regions of the dataset, ensuring the chosen medoids are both well-separated and representatives of underlying structural modes. In DIVINE, we run *k*-means NANI with *k* = 2 and use the strat_all initialization mode. Other NANI init_type variants (strat_reduced, comp_sim, div_select), as well as stochastic options (random, *k*-means++), are also supported.

#### outlier_pair

This strategy selects a structurally distinct frame (A) as an outlier based on cMSD, specifically, the frame with the lowest cMSD. It then identifies the frame (B) in the cluster that is farthest from A. Together, A and B span the approximate diameter of the cluster and serve as anchors for the split. Each frame is assigned to the closer of the two. Although simple and intuitive, this method may produce highly imbalanced splits if either A or B is an isolated outlier, resulting in a trivial subcluster.

#### splinter_split

This DIANA-inspired approach is a variant of outlier_pair, with a slightly different logic for defining the split. It first identifies an outlier frame A (as in outlier_pair) and then computes the medoid M of the remaining frames. Each frame is then assigned to the closer of the two anchors.

Due to their sensitivity to outliers, both outlier_pair and splinter_split are followed by an additional refinement step to enhance the quality and stability of the resulting partition.

### Partition Refinement

After generating an initial split using any of the anchor selection strategies, DIVINE applies a refinement step to improve cluster compactness and boundary definition. In the current implementation, this refinement is applied by default for the outlier_pair and splinter_split methods, while it is inherently integrated into the NANI-based approach through its *k*-means execution.

The refinement procedure involves performing standard *k*-means (with *k* = 2) clustering on the cluster, using the medoids of the two original subclusters as the initial seeds. This additional step helps reassign borderline conformations based on proximity to the refined medoids, helping to improve cluster boundaries and reduce unbalanced partitions.

### Clustering Quality Evaluation

DIVINE continuously tracks clustering quality metrics as a function of the number of clusters. After each split, we record the Calinski–Harabasz index (CHI) and Davies–Bouldin index (DBI) for the current partitioning level. CHI, also known as the variance ratio criterion or the pseudo-F statistic, evaluates the ratio of inter-cluster dispersion to intra-cluster compactness.^35^ Higher CHI values indicate more compact and well-separated clusters. On the other hand, DBI measures the average similarity between the most similar clusters, based on the ratio of internal spread to the distance between cluster centroids.^36^ Lower DBI values reflect higher-quality clustering. Since DIVINE builds the hierarchy in a single run, these metrics are computed and stored incrementally at each stage, yielding a complete profile of clustering quality across a range of *k*. Additionally, DIVINE generates a detailed log of the population and internal dispersion (MSD) of every cluster at each level of the hierarchy. This allows users to explicitly trace the lineage of conformational states and monitor the evolution of cluster separation and compactness as the resolution increases without the need for reruns.

To provide preliminary visual validation of our approach, we tested DIVINE on a synthetic two-dimensional benchmark dataset. The dataset was generated using a Gaussian-distributed ensemble of points (scikit-learn’s make_blobs)^20^ with sufficient standard deviation to create noisy, overlapping boundaries between some clusters (Fig. 1). This intentional overlap was designed to mimic the structural ambiguity often found in conformational ensembles, where distinct states are often bridged by transition regions. As shown in the comparisons, DIVINE (using NANI anchors) achieves visually cleaner separation compared to *k*-means NANI, particularly in resolving the decision boundaries between these closely packed groups. This improvement stems from the divisive nature of the algorithm. By isolating clusters sequentially, DIVINE prevents the position of distant centroids from influencing local decision boundaries, a common limitation in the global optimization of flat *k*-means where all centroids are updated simultaneously.

**Figure 1.**
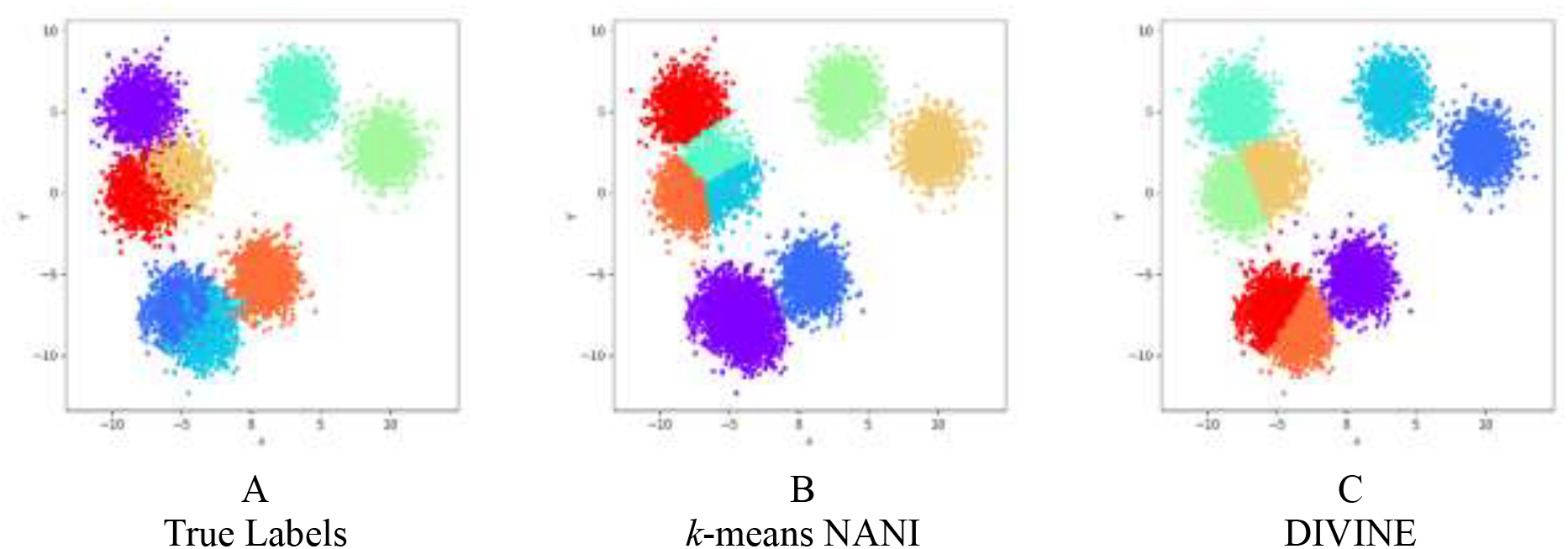
Clustering of a synthetic two-dimensional benchmark dataset. Shown are (A) the ground-truth labels for the dataset, (B) clustering obtained using flat *k*-means with NANI initialization, and (C) clustering obtained using DIVINE with NANI anchor selection.

While this simple example serves as a controlled proof of concept, we emphasize that it does not reflect the complexity of molecular dynamics data; therefore, we proceed to a rigorous validation using a biomolecular system.

## SYSTEM AND PROTOCOLS

### HP35 Folding Trajectory

Our primary test system is a 305 μs molecular dynamics simulation of a fast-folding villin headpiece subdomain variant (Nle/Nle double mutant), performed at 360 K by D. E. Shaw Research.^37^ The trajectory comprises approximately 1.5 million frames saved every 200 ps. This system is a 35-residue protein featuring three helical segments: helix 1 spans residues 4–10, helix 2 includes residues 15–19, and helix 3 covers residues 23–32.^38^ To prepare the data, we discarded all frames prior to the 5,000^th^ frame to exclude initial equilibration, and aligned the remaining frames to the 5,000^th^ frame. For structural comparison, we selected the backbone atoms following prior studies: the N atom of residue 1; the N, Cα, and C atoms of residues 2 through 34; and the N atom of residue 35. To reduce data volume, we applied a 10-frame sieving interval, giving us a final set of 151,105 frames for clustering.

To contextualize DIVINE’s performance, we also applied scikit-learn’s BKM algorithm, on the same HP35 dataset. We tested two cluster selection strategies: biggest inertia and largest cluster as well as two initialization settings, random and *k*-means++. As noted earlier, because scikit-learn’s BKM does not natively preserve the hierarchy of splits, we had to run the algorithm for each desired number of clusters (with 3 independent runs for each setting to gauge variability). This allowed us to compare the internal metrics with that of DIVINE.

## RESULTS AND DISCUSSION

### Cluster Selection Strategies

Since each iteration in DIVINE begins by selecting a cluster to split, we first evaluated different strategies for making this choice. DIANA selects the cluster with the largest diameter (i.e., the most dissimilar pair of points), but as discussed earlier, this approach is highly sensitive to outliers and can lead to fragmented or structurally uninformative splits. BKM uses either the largest cluster (by size) or the one with the biggest inertia (sum of squared errors). While this reduces direct sensitivity to individual outliers, it still introduces a bias toward larger clusters and may overlook smaller, yet heterogeneous, clusters.

To address these limitations, we explored metrics that capture dispersion more explicitly while remaining computationally efficient. We first implemented two measures: MSD and radius, which can be evaluated in linear time with respect to cluster size. MSD estimates the average pairwise squared distance, while radius measures the maximum distance from the cluster medoid to any member. Both can identify clusters with high internal variance. However, our initial tests revealed a consistent pattern: MSD and radius frequently prioritized dispersed and noisy clusters early in the hierarchy. As shown in Table S1, these splits produced long tails of high-MSD small clusters branching from a dominant supercluster.

To reduce this bias, we introduced the weighted_MSD criterion, which multiplies the cluster’s MSD by its size. This adjustment steers selection toward large, heterogeneous clusters rather than small, high-variance ones. As a result, the early stages of the hierarchy exhibit more balanced cluster populations and fewer trivial outlier branches.

In Fig. 2, we compare the performance of these selection strategies using both CHI and DBI. Both MSD and radius, with the largest_cluster variant of BKM consistently yielded lower CHI and higher DBI scores, indicating less compact and less separated clusters. In contrast, weighted_MSD outperformed these methods and matched or exceeded BKM’s biggest_inertia method across the full range of k. Error bars are shown only for BKM, as its reliance on random initialization leads to run-to-run variability. Based on these observations, we adopted weighted_MSD as the default splitting criterion for DIVINE and biggest_inertia for BKM to be used for further analyses.

**Figure 2.**
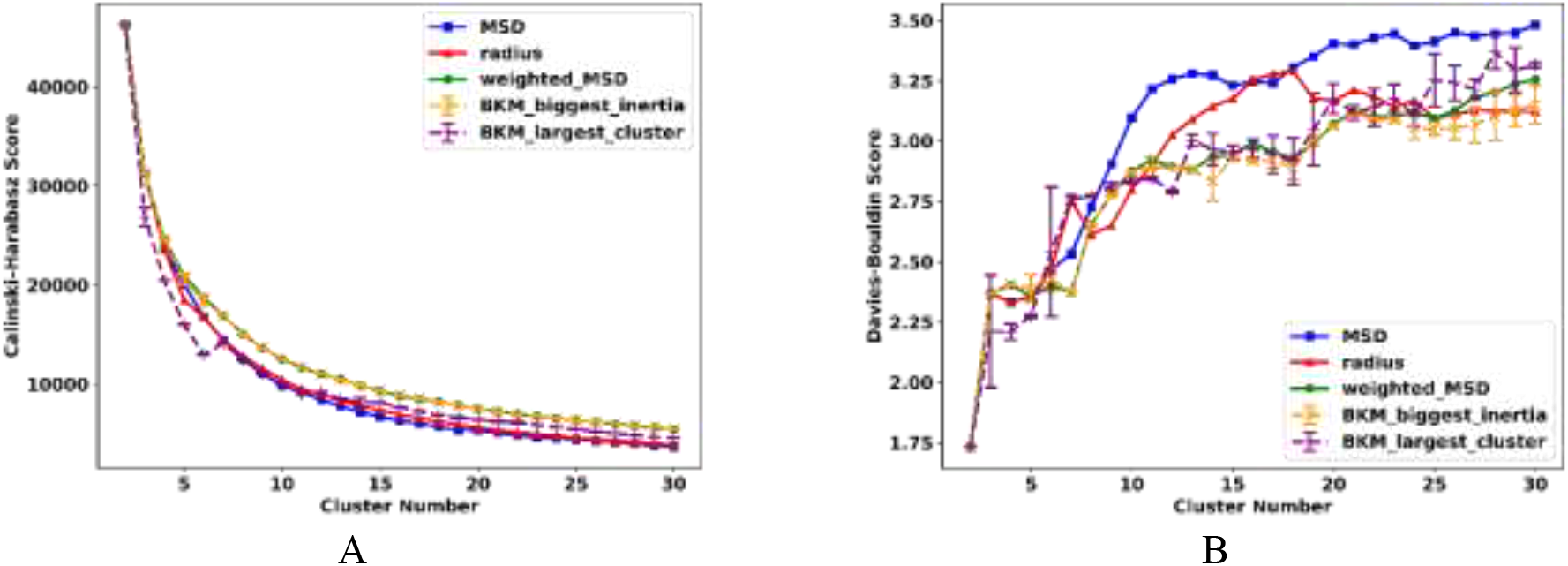
Change in (A) Calinski-Harabasz (higher is better) and (B) Davies-Bouldin (lower is better) indices for HP35 after DIVINE clustering from *k* = 1 to 30, comparing performance across different cluster selection strategies with NANI anchor selection.

In addition to choosing a good splitting metric, we also found it useful to require a minimum split size to guard against trivial splits. As described in Theory, we introduced a threshold parameter requiring that both resulting subclusters contain at least a minimum fraction of the total frames, or else that split is rejected, and the next top cluster is chosen for splitting. However, we note that an aggressive threshold can eventually prevent any split when cluster sizes become naturally small. In fact, with a 5% threshold, DIVINE could only split up to about 10 clusters on HP35 before all remaining splits would violate the size rule. Interestingly, under that condition, the three split criteria all led to the same clusters when *k* = 7 (Table S2).

In summary, a moderate threshold can improve cluster balance, but it should be set based on the intended number of clusters. For all main runs, we disabled this threshold criterion unless otherwise noted, to allow the hierarchy to fully emerge for screening purposes.

### Anchor Strategies and Refinement

After selecting a cluster to split, DIVINE determines the anchor frames that define the initial subdivision. These anchors serve as seeds that represent the emerging subclusters, and their selection strongly influences the quality and shape of the resulting hierarchy. DIVINE provides three deterministic anchor strategies: NANI, outlier_pair, and splinter_split, each reflecting a different principle of cluster initialization.

In Fig. 3, we report the CHI and DBI values obtained from each strategy across a range of *k* = 1 to 30. Based on DBI alone, splinter_split appears favorable, especially at low *k*, suggesting the formation of compact, well-separated clusters. However, CHI reveals a different trend: splinter_split consistently underperforms, with flat CHI scores across the screening range, indicating limited improvement in between-cluster separation.

**Figure 3.**
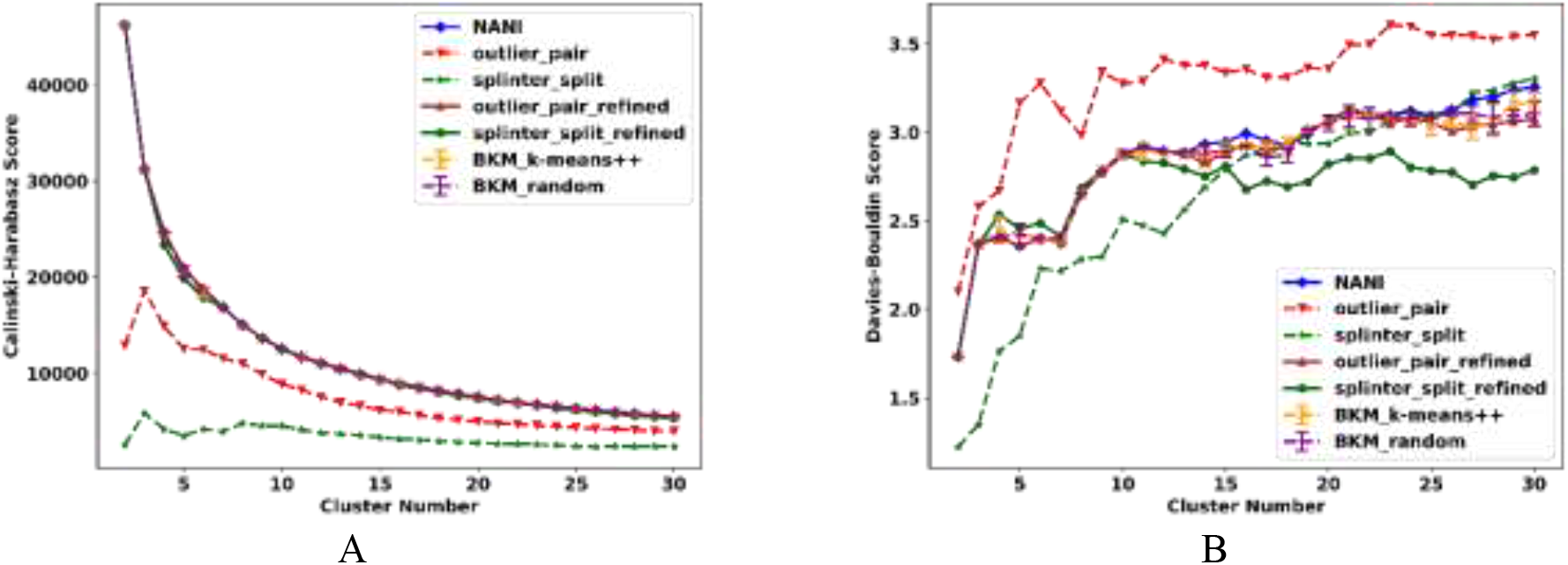
Change in (A) Calinski-Harabasz (higher is better) and (B) Davies-Bouldin (lower is better) indices for HP35 after DIVINE clustering from *k* = 1 to 30, comparing performance across different anchor methods.

To explain this discrepancy, we examined the cluster populations and MSDs at *k* = 7 (Table 1). The results showed that splinter_split produced a dominant cluster containing over 94% of the frames, while the remaining clusters accounted for less than 2% each. These small clusters typically exhibited high internal MSD, suggesting that they were isolated based on outlier frames rather than structural significance. A similar trend was observed for outlier_pair, although the imbalance was slightly less extreme. The cumulative distribution plots in Fig. S3 further visualize this disparity, contrasting the sharp population skew of the unrefined methods against the more balanced distribution achieved by NANI.

**Table 1.** Cluster populations and MSDs of each cluster at *k* = 7 from DIVINE runs using NANI, outlier_pair, and splinter_split as anchors.

| NANI |  |  | outlier_pair |  |  | splinter_split |  |  |
| --- | --- | --- | --- | --- | --- | --- | --- | --- |
| Population | Fraction | MSD | Population | Fraction | MSD | Population | Fraction | MSD |
| 6347 | 4.20 | 88.16 | 3299 | 2.18 | 91.40 | 345 | 0.23 | 84.54 |
| 7681 | 5.08 | 87.81 | 3590 | 2.38 | 78.97 | 541 | 0.36 | 92.72 |
| 10569 | 6.99 | 38.73 | 12732 | 8.43 | 95.49 | 826 | 0.55 | 86.96 |
| 11774 | 7.79 | 80.73 | 13318 | 8.81 | 99.14 | 1066 | 0.71 | 71.21 |
| 13401 | 8.87 | 74.74 | 14661 | 9.70 | 58.40 | 2509 | 1.66 | 84.04 |
| 36677 | 24.27 | 16.89 | 17436 | 11.54 | 22.33 | 2690 | 1.78 | 88.81 |
| 64656 | 42.79 | 9.30 | 86069 | 56.96 | 13.33 | 143128 | 94.72 | 43.89 |

To mitigate this, we applied a refinement step for the outlier-based anchors. After the initial outlier_pair or splinter_split division, we performed a *k*-means (with *k* = 2) refinement using the medoids as initial centers (see Theory). These refined versions (shown as outlier_pair_refined and splinter_split_refined in the plots) yielded significantly better results. As shown in Fig. 3, both refined anchors show marked CHI gains, especially at moderate *k*, and maintain or improve DBI scores that match NANI and the BKM variants. Table 2 confirms improved population balance: the largest cluster drops below 50%, and several secondary clusters grow beyond 5%, making the resulting clusters more interpretable.

**Table 2.** Cluster populations and MSDs of each cluster at *k* = 7 from DIVINE runs using the NANI and the refined variants of outlier_pair and splinter_split as anchors.

| NANI |  |  | outlier_pair_refined |  |  | splinter_split_refined |  |  |
| --- | --- | --- | --- | --- | --- | --- | --- | --- |
| Population | Fraction | MSD | Population | Fraction | MSD | Population | Fraction | MSD |
| 6347 | 4.20 | 88.16 | 6372 | 4.22 | 88.35 | 6371 | 4.22 | 87.98 |
| 7681 | 5.08 | 87.81 | 7667 | 5.07 | 87.77 | 7662 | 5.07 | 88.10 |
| 10569 | 6.99 | 38.73 | 10571 | 7.00 | 38.75 | 11796 | 7.81 | 80.72 |
| 11774 | 7.79 | 80.73 | 11745 | 7.77 | 80.64 | 13354 | 8.84 | 74.63 |
| 13401 | 8.87 | 74.74 | 13405 | 8.87 | 74.76 | 13403 | 8.87 | 40.29 |
| 36677 | 24.27 | 16.89 | 36688 | 24.28 | 16.90 | 32312 | 21.38 | 15.45 |
| 64656 | 42.79 | 9.30 | 64657 | 42.79 | 9.30 | 66207 | 43.82 | 8.94 |

In contrast, the NANI strategy yielded structurally significant partitions without the need for refinement. While we do not impose an artificial requirement for “balanced” cluster sizes, NANI avoids the severe imbalance observed in unrefined methods, where the algorithm repeatedly isolates small, dispersed groups, while leaving the bulk of the conformational landscape unresolved. At *k* = 7, the largest cluster contained ∼43% of frames, with the remainder distributed among clusters ranging from ∼4% to ∼24%. These ratios fall within the ranges reported in previous studies^13,18^ and support our choice of NANI as the default anchor strategy in DIVINE.

### Cluster Number Selection

Beyond evaluating clustering quality across anchor strategies, a central motivation for divisive algorithms like DIVINE lies in their inherent ability to facilitate cluster screening. By constructing the full hierarchy in a single run, DIVINE enables users to identify optimal clustering resolutions, defined here as the values of *k* corresponding to extrema or significant inflection points in the validity metrics, without requiring separate runs for each target *k*. To assess this capability, we evaluated how well each anchor strategy supports the selection of *k*, not only by examining absolute metric performance, but also by analyzing how reliably they indicate where to partition the tree into meaningful clusters.

Our approach involved two levels of analysis. First, we identified global optima, the values of *k* at which CHI reaches its maximum or DBI reaches its minimum within the screened range. These points represent the optimal number of clusters according to each metric. Second, we examined local optima by computing the second finite-difference derivatives of the CHI and DBI curves. Distinct peaks or dips in these second-derivative profiles highlight regions where splitting becomes more stable, and where further increases in splits contribute less to overall improvement. Because these metrics often bias toward trivial partitions (e.g., *k* = 1, 2, or 3), we restricted our search for optima to the range *k* ≥ 5, which corresponds to the minimum resolution at which multiple states emerge for this system. For other systems, this lower bound can be adjusted based on the expected number of dominant basins, with the same principle of excluding trivial partitions applied generally. These inflection points can reveal physically significant partitions (or optimal cluster counts), even if they do not correspond to the global extremum.

Table 3 summarizes the optimal number of clusters based on these metrics. All DIVINE strategies showed global CHI maxima at *k* = 5 and secondary peaks at *k* = 6, suggesting consistent identification of major conformational states. For DBI, both NANI and outlier_pair reached minima at *k* = 5, while splinter_split achieved its minimum at *k* = 7, reflecting its preference for finer-grained partitioning. Second-derivative trends were more revealing: all DIVINE anchors showed inflection points at *k* = 7. Detailed profiles of these metrics and their derivatives for each anchor strategy are provided in Fig. S4-S6. Moreover, to verify that these features are robust to sampling noise, we performed a consistency analysis using frame sieving intervals ranging from 1 to 100 with NANI as anchor (Fig. S8). The optimal cluster count indicated the maximum second derivative of DBI remained consistent across these sampling densities, confirming that the identified minima reflect stable structural features rather than sampling artifacts.

**Table 3.** Preferred numbers of clusters for HP35, identified using global and local optima from CHI and DBI metrics following clustering.

| **metric\anchor** | **DIVINE**<br>**NANI** | **DIVINE**<br>**outlier_pair** | **DIVINE**<br>**splinter_split** | **BKM\***<br>**k-means++** | **BKM\***<br>**random** |
| --- | --- | --- | --- | --- | --- |
| Max. CH | 5 | 5 | 5 | 5/5/5 | 5/5/5 |
| 2nd Max. CH | 6 | 6 | 6 | 6/6/6 | 6/6/6 |
| Min. 2nd Der. CH | - | - | - | - | - |
| 2nd Min. 2nd Der. CH | - | - | - | - | - |
| Min. DBI | 5 | 5 | 7 | 5/7/5 | 5/7/5 |
| 2nd Min. DBI | 7 | 7 | 5 | 7/6/7 | 6/6/6 |
| Max 2nd Der. DBI | 7 | 7 | 7 | 7/28/7 | 22/7/29 |
| 2nd Max 2nd Der. DBI | 18 | 14 | 16 | 14/7/27 | 20/18/17 |

\*BKM results shown as triplicate runs ( $k$ -means++ and random initializations).

In contrast, BKM strategies exhibited scattered and inconsistent inflection points across runs (Fig. S7), highlighting the challenges of reproducible model selection under stochastic initialization.

Based on this analysis, we selected *k* = 7 for structural investigation. Fig. 4 shows the overlaps of the best representative frames from each of the seven clusters identified by DIVINE using NANI anchors and weighted_MSD, capturing distinct structural motifs within the HP35 folding trajectory. These same states were identified in previous studies using the *k*-means NANI clustering of the system,^13^ highlighting consistency across methods. Moreover, since both algorithms successfully recover the same key conformational states, it is instructive to clarify their distinct roles in the analysis pipeline. While *k*-means NANI performs a global optimization that refines all centroids simultaneously, it requires independent execution for each target *k*, meaning that clusters found at *k* = 6 have no guaranteed mathematical relationship to those at *k* = 7. In contrast, DIVINE’s primary advantage lies in its sequential top-down construction: it generates the entire clustering hierarchy in a single pass, where each step naturally yields the next cluster count (*k* → *k* + 1), while preserving the parent-child lineage of split clusters. This makes DIVINE the preferred choice for exploring the hierarchical connectivity of conformational states (e.g., observing how the unfolded basin subdivides), whereas *k*-means NANI remains the standard for producing a flat global partition once a specific *k* is selected.

**Figure 4.**
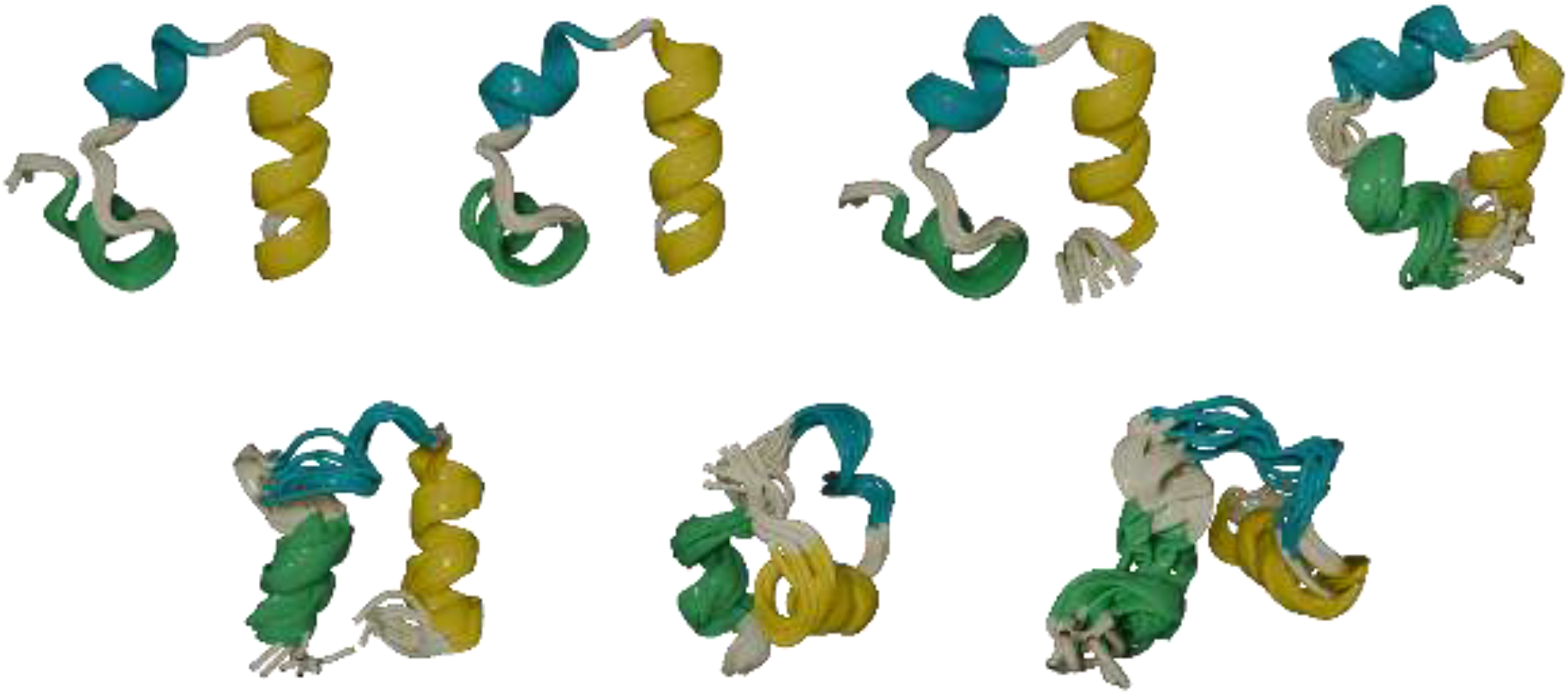
Overlaps of best representative structures from the seven clusters of HP35 after performing DIVINE using NANI as anchors and weighted_MSD as cluster selection strategy. Helices 1, 2, and 3 are colored green, cyan, and yellow, respectively. Each overlay consists of the cluster medoid with the 10 frames having the highest similarity to that medoid (total of 11 structures per cluster).

### Transferability and Practical Recommendations

While our benchmarks focused on the HP35 folding trajectory, a system chosen for its extensive sampling of diverse conformational states ranging from unfolded, intermediate, to native structures, we recognize that different molecular systems (e.g., nucleic acids or large multi-domain complexes) must exhibit distinct density profiles. However, the performance trends we observed are rooted in geometric principles rather than system-specific features. For instance, we see the tendency of unweighted variance metrics like MSD or radius to “shave off” outliers one by one. By penalizing small clusters, the weighted variance criterion effectively mitigates this fragmentation, a behavior we expect to be transferable to other macromolecular datasets.

For users applying DIVINE to novel systems, we recommend the following to select the optimal parameters. First, we advise starting with the default settings (weighted_MSD split criterion and NANI anchors), as this combination offers the most robust balance between cluster population and structural compactness for general purpose analysis. Second, the users should leverage DIVINE’s single-pass capability to inspect resulting CHI and DBI scores; a lack of distinct peaks or plateaus in these metrics may indicate that the split criterion may not align with the system’s specific conformational landscape. Finally, if the analytical goal of the user is to specifically identify rare high-variance states (e.g., cryptic pockets or transition states) rather than dominant basins, they may benefit from switching the criterion to radius or MSD. Crucially, because DIVINE is highly efficient, as demonstrated in the following subsection, it can screen millions of frames in minutes. Performing these exploratory reruns with alternative settings is computationally affordable and does not create a significant bottleneck.

### Scalability and Performance

To evaluate full runtime performance, we benchmarked all methods on two datasets (Table 4): a 150k frame subset and the full 1.5 million-frame trajectory of HP35. On the smaller dataset, all DIVINE anchor variants were completed in less than 40 seconds. In comparison, BKM with *k*-means++ and random initialization took 205.47 and 139.95 seconds, respectively, which are approximately four times longer compared to the DIVINE methods.

**Table 4:** Runtime (s) to complete a DIVINE screening (*k* = 30) with each anchor method.

| anchor | ~151k frames (s) | ~1.5m frames (s) |
| --- | --- | --- |
| NANI | 34.75 | 283.38 |
| outlier_pair | 33.39 | 312.06 |
| splinter_split | 37.29 | 559.25 |
| BKM_$k$-means++ | 205.47 | 1418.26 |
| BKM_random | 139.95 | 1324.40 |

As we screen the full 1.5 million HP35 trajectory, the differences become even more significant. NANI and outlier_pair anchors were completed in around six minutes. Even the more computationally demanding splinter_split anchor finished in under 10 minutes. In contrast, BKM required over 23 minutes (1418.26 seconds) using *k*-means++ initialization, and 22 minutes (1324.40 seconds) for the random variant. However, BKM’s slower performance is partly due to implementation limitations. As mentioned, standard libraries like scikit-learn do not expose the internal clustering hierarchy, so unless the actual code is modified, the entire process must be repeated for each target value of *k*. While this is a technical detail, it highlights one of the advantages of DIVINE: it generates and retains all partitions in a single pass, enabling fast and reproducible screening even at large scales.

## CONCLUSIONS

We presented DIVINE, a deterministic, top-down clustering algorithm for MD trajectory analysis. DIVINE builds a complete, reproducible clustering hierarchy in a single pass by iteratively splitting clusters using cluster-wise metrics and robust anchor selection. Unlike conventional methods, DIVINE avoids pairwise distance matrices, random seeds, and repeated runs, enabling rapid and memory-efficient exploration of the conformational landscape.

The algorithm selects clusters for division based on dispersion and size-driven criteria, including MSD, radius, and weighted MSD, and supports multiple anchor strategies: NANI-based initialization, outlier-pair anchoring, and a DIANA-inspired splintering scheme. More importantly, DIVINE records quality metrics at each step of the hierarchy, enabling users to evaluate clustering performance globally across a range of cluster counts without rerunning the algorithm.

Our benchmarks on the HP35 trajectory demonstrate that DIVINE matches or surpasses bisecting *k*-means (with *k*-means++) in clustering quality, while significantly reducing runtime and eliminating variability across runs. We were able to complete the screening of the full 1.5 million HP35 frames in under six minutes on a single CPU core on a standard workstation, while producing well-separated and balanced clusters. The default combination of weighted MSD for cluster selection and NANI-based anchors consistently yielded stable partitions that reflected meaningful and previously reported structural states.

Finally, we address the role of dimensionality reduction techniques such as Principal Component Analysis (PCA) or Time-lagged Independent Component Analysis (TICA), which are frequently employed to preprocess high-dimensional data prior to clustering. While these methods effectively reduce computational overhead and can help filter noise, they necessarily project the ensemble onto a lower-dimensional representation. Such compression can risk obscuring subtle yet chemically relevant structural differences, particularly when such differences do not align with the dominant variance or slow collective modes.^16,39^ A key advantage of DIVINE is that its computational efficiency allows dimensionality reduction to be used as an option rather than a necessity, depending on the user’s analytical goals.

Methodologically, DIVINE’s core strength lies in its divisive, top-down construction of the hierarchy. Unlike agglomerative or partitional methods, which are limited to either bottom-up heuristic or flat partitioning, this divisive strategy enables the global structural information to guide each recursive splitting. This allows a clearer representation of the system’s conformational landscape, particularly in cases where dominant modes span large portions in the trajectory. This also enables DIVINE to leverage global structural information before committing to local partitions, allowing for a more coherent and chemically meaningful decomposition of an MD trajectory.

## SUPPORTING INFORMATION

Tables with the cluster statistics and figures displaying structural overlaps, validity metric profiles, and robustness analyses (PDF).

## ACKNOWLEDGEMENTS

RAMQ, LC, and JBWS thank support from the National Institute of General Medical Sciences of the National Institutes of Health under award number R35GM150620.

## AUTHOR CONTRIBUTIONS

JBWS: Data curation; Formal analysis; Investigation; Software; Validation; Visualization; Writing.

LC: Data curation; Formal analysis; Investigation; Software; Validation; Visualization; Writing.

RAMQ: Formal analysis; Methodology; Conceptualization; Investigation; Software; Writing; Funding acquisition; Supervision; Resources.

## DATA AND SOFTWARE AVAILABILITY

The code for DIVINE can be found here: https://github.com/mqcomplab/MDANCE.

## TOC Graphic

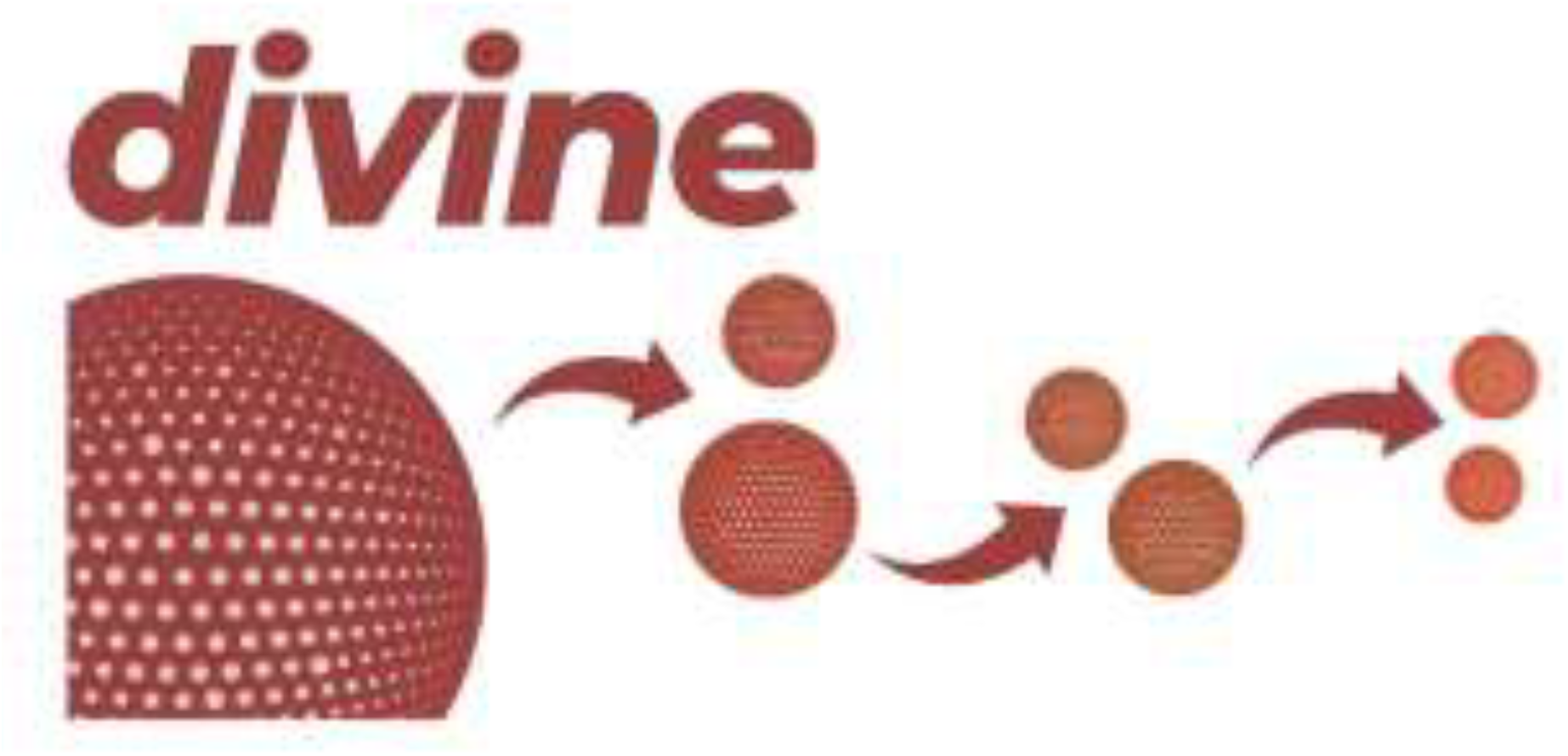

## REFERENCES

(1) Karplus, M.; McCammon, J. A. Molecular Dynamics Simulations of Biomolecules. Nat. Struct Biol. 2002, 9 (9), 646–652. 10.1038/nsb0902-646.

(2) Arnittali, M.; Rissanou, A. N.; Harmandaris, V. Structure Of Biomolecules Through Molecular Dynamics Simulations. Procedia Computer Science 2019, 156, 69–78. 10.1016/j.procs.2019.08.181.

(3) Pierce, L. C. T.; Salomon-Ferrer, R.; Augusto F. De Oliveira, C.; McCammon, J. A.; Walker, R. C. Routine Access to Millisecond Time Scale Events with Accelerated Molecular Dynamics. J. Chem. Theory Comput. 2012, 8 (9), 2997–3002. 10.1021/ct300284c.

(4) Shaw, D. E.; Deneroff, M. M.; Dror, R. O.; Kuskin, J. S.; Larson, R. H.; Salmon, J. K.; Young, C.; Batson, B.; Bowers, K. J.; Chao, J. C.; Eastwood, M. P.; Gagliardo, J.; Grossman, J. P.; Ho, C. R.; Ierardi, D. J.; Kolossváry, I.; Klepeis, J. L.; Layman, T.; McLeavey, C.; Moraes, M. A.; Mueller, R.; Priest, E. C.; Shan, Y.; Spengler, J.; Theobald, M.; Towles, B.; Wang, S. C. Anton, a Special-Purpose Machine for Molecular Dynamics Simulation. Commun. ACM 2008, 51 (7), 91–97. 10.1145/1364782.1364802.

(5) Harvey, M. J.; Giupponi, G.; Fabritiis, G. D. ACEMD: Accelerating Biomolecular Dynamics in the Microsecond Time Scale. J. Chem. Theory Comput. 2009, 5 (6), 1632–1639. 10.1021/ct9000685.

(6) De Paris, R.; Quevedo, C. V.; Ruiz, D. D.; Norberto De Souza, O.; Barros, R. C. Clustering Molecular Dynamics Trajectories for Optimizing Docking Experiments. Computational Intelligence and Neuroscience 2015, 2015, 1–9. 10.1155/2015/916240.

(7) Shao, J.; Tanner, S. W.; Thompson, N.; Cheatham, T. E. Clustering Molecular Dynamics Trajectories: 1. Characterizing the Performance of Different Clustering Algorithms. J. Chem. Theory Comput. 2007, 3 (6), 2312–2334. 10.1021/ct700119m.

(8) Keller, B.; Daura, X.; Van Gunsteren, W. F. Comparing Geometric and Kinetic Cluster Algorithms for Molecular Simulation Data. The Journal of Chemical Physics 2010, 132 (7), 074110. 10.1063/1.3301140.

(9) Salsbury Jr, F. R. Molecular Dynamics Simulations of Protein Dynamics and Their Relevance to Drug Discovery. Current Opinion in Pharmacology 2010, 10 (6), 738–744. 10.1016/j.coph.2010.09.016.

(10) Knaggs, M. H.; Salsbury, F. R.; Edgell, M. H.; Fetrow, J. S. Insights into Correlated Motions and Long-Range Interactions in CheY Derived from Molecular Dynamics Simulations. Biophysical Journal 2007, 92 (6), 2062–2079. 10.1529/biophysj.106.081950.

(11) Ikotun, A. M.; Ezugwu, A. E.; Abualigah, L.; Abuhaija, B.; Heming, J. K-Means Clustering Algorithms: A Comprehensive Review, Variants Analysis, and Advances in the Era of Big Data. Information Sciences 2023, 622, 178–210. 10.1016/j.ins.2022.11.139.

(12) Arthur, D.; Vassilvitskii, S. K-Means++: The Advantages of Careful Seeding. In Proceedings of the Eighteenth Annual ACM-SIAM Symposium on Discrete Algorithms; SODA ‘07; Society for Industrial and Applied Mathematics: USA, 2007; pp 1027–1035.

(13) Chen, L.; Roe, D. R.; Kochert, M.; Simmerling, C.; Miranda-Quintana, R. A. K-Means NANI: An Improved Clustering Algorithm for Molecular Dynamics Simulations. J. Chem. Theory Comput. 2024, 20 (13), 5583–5597. 10.1021/acs.jctc.4c00308.

(14) Sculley, D. Web-Scale k-Means Clustering. In Proceedings of the 19th international conference on World wide web; ACM: Raleigh North Carolina USA, 2010; pp 1177–1178. 10.1145/1772690.1772862.

(15) Feizollah, A.; Anuar, N. B.; Salleh, R.; Amalina, F. Comparative Study of K-Means and Mini Batch k-Means Clustering Algorithms in Android Malware Detection Using Network Traffic Analysis. In 2014 International Symposium on Biometrics and Security Technologies (ISBAST); IEEE: Kuala Lumpur, Malaysia, 2014; pp 193–197. 10.1109/ISBAST.2014.7013120.

(16) Scherer, M. K.; Trendelkamp-Schroer, B.; Paul, F.; Pérez-Hernández, G.; Hoffmann, M.; Plattner, N.; Wehmeyer, C.; Prinz, J.-H.; Noé, F. PyEMMA 2: A Software Package for Estimation, Validation, and Analysis of Markov Models. J. Chem. Theory Comput. 2015, 11 (11), 5525–5542. 10.1021/acs.jctc.5b00743.

(17) Xu, D.; Tian, Y. A Comprehensive Survey of Clustering Algorithms. Ann. Data. Sci. 2015, 2 (2), 165–193. 10.1007/s40745-015-0040-1.

(18) Chen, L.; Brylle Woody Santos, J.; Gaza, J.; Perez, A.; Miranda-Quintana, R. A. Hierarchical Extended Linkage Method (HELM)’s Deep Dive into Hybrid Clustering Strategies. J. Chem. Inf. Model. 2025. 10.1021/acs.jcim.5c00539.

(19) Wang, Y.; Moseley, B. An Objective for Hierarchical Clustering in Euclidean Space and Its Connection to Bisecting K-Means. AAAI 2020, 34 (04), 6307–6314. 10.1609/aaai.v34i04.6099.

(20) Pedregosa, F.; Varoquaux, G.; Gramfort, A.; Michel, V.; Thirion, B.; Grisel, O.; Blondel, M.; Prettenhofer, P.; Weiss, R.; Dubourg, V.; Vanderplas, J.; Passos, A.; Cournapeau, D.; Brucher, M.; Perrot, M.; Duchesnay, E. Scikit-Learn: Machine Learning in Python. Journal of Machine Learning Research 2011, 12, 2825–2830.

(21) Karypis, M. S. G.; Kumar, V.; Steinbach, M. A Comparison of Document Clustering Techniques. In TextMining Workshop at KDD2000 (May 2000); 2000; pp 428–439.

(22) Kaufman, L.; Rousseeuw, P. J. Finding Groups in Data: An Introduction to Cluster Analysis, 1st ed.; Wiley Series in Probability and Statistics; Wiley, 1990. 10.1002/9780470316801.

(23) Sharf, Z.; Razzak, M. The Informative Vector Selection in Active Learning Using Divisive Analysis. International Journal of Advanced Computer Science and Applications 2017, 8 (10).

(24) Miranda-Quintana, R. A.; Bajusz, D.; Rácz, A.; Héberger, K. Extended Similarity Indices: The Benefits of Comparing More than Two Objects Simultaneously. Part 1: Theory and Characteristics†. J Cheminform 2021, 13 (1), 32. 10.1186/s13321-021-00505-3.

(25) Miranda-Quintana, R. A.; Rácz, A.; Bajusz, D.; Héberger, K. Extended Similarity Indices: The Benefits of Comparing More than Two Objects Simultaneously. Part 2: Speed, Consistency, Diversity Selection. J Cheminform 2021, 13 (1), 33. 10.1186/s13321-021-00504-4.

(26) Chang, L.; Perez, A.; Miranda-Quintana, R. A. Improving the Analysis of Biological Ensembles through Extended Similarity Measures. Phys. Chem. Chem. Phys. 2022, 24 (1), 444–451. 10.1039/D1CP04019G.

(27) Chen, L.; Leung, J. M. G.; Zsigmond, K.; Chong, L. T.; Miranda-Quintana, R. A. SHINE: Deterministic Many-to-Many Clustering of Molecular Pathways. J. Chem. Inf. Model. 2025, 65 (10), 4775–4782. 10.1021/acs.jcim.5c00240.

(28) Chen, L.; Smith, M.; Roe, D. R.; Miranda-Quintana, R. A. Extended Quality (eQual): Radial Threshold Clustering Based on n -Ary Similarity. J. Chem. Inf. Model. 2025, 65 (10), 5062–5070. 10.1021/acs.jcim.4c02341.

(29) López-Pérez, K.; Kim, T. D.; Miranda-Quintana, R. A. iSIM: Instant Similarity. Digital Discovery 2024, 3 (6), 1160–1171. 10.1039/D4DD00041B.

(30) Rácz, A.; Mihalovits, L. M.; Bajusz, D.; Héberger, K.; Miranda-Quintana, R. A. Molecular Dynamics Simulations and Diversity Selection by Extended Continuous Similarity Indices. J. Chem. Inf. Model. 2022, 62 (14), 3415–3425. 10.1021/acs.jcim.2c00433.

(31) Rácz, A.; Dunn, T. B.; Bajusz, D.; Kim, T. D.; Miranda-Quintana, R. A.; Héberger, K. Extended Continuous Similarity Indices: Theory and Application for QSAR Descriptor Selection. J Comput Aided Mol Des 2022, 36 (3), 157–173. 10.1007/s10822-022-00444-7.

(32) Chen, L.; Roe, D. R.; Miranda-Quintana, R. A. CADENCE: Clustering Algorithm—Density-Based Exploration and Novelty Clustering with Efficiency. J. Chem. Inf. Model. 2025, acs.jcim.5c00392. 10.1021/acs.jcim.5c00392.

(33) Pérez, K. L.; Jung, V.; Chen, L.; Huddleston, K.; Miranda-Quintana, R. A. BitBIRCH: Efficient Clustering of Large Molecular Libraries. Digital Discovery 2025, 4 (4), 1042–1051. 10.1039/D5DD00030K.

(34) Chen, L.; Mondal, A.; Perez, A.; Miranda-Quintana, R. A. Protein Retrieval via Integrative Molecular Ensembles (PRIME) through Extended Similarity Indices. J. Chem. Theory Comput. 2024, 20 (14), 6303–6315. 10.1021/acs.jctc.4c00362.

(35) Calinski, T.; Harabasz, J. A Dendrite Method for Cluster Analysis. Comm. in Stats. - Theory & Methods 1974, 3 (1), 1–27. 10.1080/03610927408827101.

(36) Davies, D. L.; Bouldin, D. W. A Cluster Separation Measure. IEEE Trans. Pattern Anal. Mach. Intell. 1979, PAMI-1 (2), 224–227. 10.1109/TPAMI.1979.4766909.

(37) Piana, S.; Lindorff-Larsen, K.; Shaw, D. E. Protein Folding Kinetics and Thermodynamics from Atomistic Simulation. Proc. Natl. Acad. Sci. U.S.A. 2012, 109 (44), 17845–17850. 10.1073/pnas.1201811109.

(38) Klem, H.; Hocky, G. M.; McCullagh, M. Size-and-Shape Space Gaussian Mixture Models for Structural Clustering of Molecular Dynamics Trajectories. J. Chem. Theory Comput. 2022, 18 (5), 3218–3230. 10.1021/acs.jctc.1c01290.

(39) Husic, B. E.; Pande, V. S. Markov State Models: From an Art to a Science. J. Am. Chem. Soc. 2018, 140 (7), 2386–2396. 10.1021/jacs.7b12191.

