## Supplementary Information for "Divide and Cluster: The DIVINE Framework for Deterministic Top-Down Analysis of Molecular Dynamics Trajectories"

### SUPPORTING INFORMATION

**Table S1.** Cluster populations and MSDs of each cluster at $k = 7$ from DIVINE runs using NANI anchors and various cluster selection strategies, without applying a threshold.

| MSD | | | radius | | | weighted_MSD | | |
|---|---|---|---|---|---|---|---|---|
| **Population** | **Fraction** | **MSD** | **Population** | **Fraction** | **MSD** | **Population** | **Fraction** | **MSD** |
| 2632 | 1.74 | 86.75 | 2632 | 1.74 | 86.75 | 6347 | 4.20 | 88.16 |
| 3715 | 2.46 | 68.79 | 3715 | 2.46 | 68.79 | 7681 | 5.08 | 87.81 |
| 3742 | 2.48 | 76.77 | 5827 | 3.86 | 73.82 | 10569 | 6.99 | 38.73 |
| 3939 | 2.61 | 82.69 | 5947 | 3.94 | 75.43 | 11774 | 7.79 | 80.73 |
| 11774 | 7.79 | 80.73 | 7681 | 5.08 | 87.81 | 13401 | 8.87 | 74.74 |
| 13401 | 8.87 | 74.74 | 13401 | 8.87 | 74.74 | 36677 | 24.27 | 16.89 |
| 111902 | 74.06 | 18.46 | 111902 | 74.06 | 18.46 | 64656 | 42.79 | 9.30 |

**Table S2.** Cluster populations and MSDs of each cluster at $k = 7$ from DIVINE runs using NANI anchors and various cluster selection strategies, with a threshold of 5%.

| MSD | | | radius | | | weighted_MSD | | |
|---|---|---|---|---|---|---|---|---|
| **Population** | **Fraction** | **MSD** | **Population** | **Fraction** | **MSD** | **Population** | **Fraction** | **MSD** |
| 7681 | 5.08 | 87.81 | 7681 | 5.08 | 87.81 | 7681 | 5.08 | 87.81 |
| 10569 | 6.99 | 38.73 | 10569 | 6.99 | 38.73 | 10569 | 6.99 | 38.73 |
| 13401 | 8.87 | 74.74 | 13401 | 8.87 | 74.74 | 13401 | 8.87 | 74.74 |
| 18121 | 11.99 | 95.13 | 18121 | 11.99 | 95.13 | 18121 | 11.99 | 95.13 |
| 23042 | 15.25 | 9.13 | 23042 | 15.25 | 9.13 | 23042 | 15.25 | 9.13 |
| 36677 | 24.27 | 16.89 | 36677 | 24.27 | 16.89 | 36677 | 24.27 | 16.89 |
| 41614 | 27.54 | 7.57 | 41614 | 27.54 | 7.57 | 41614 | 27.54 | 7.57 |

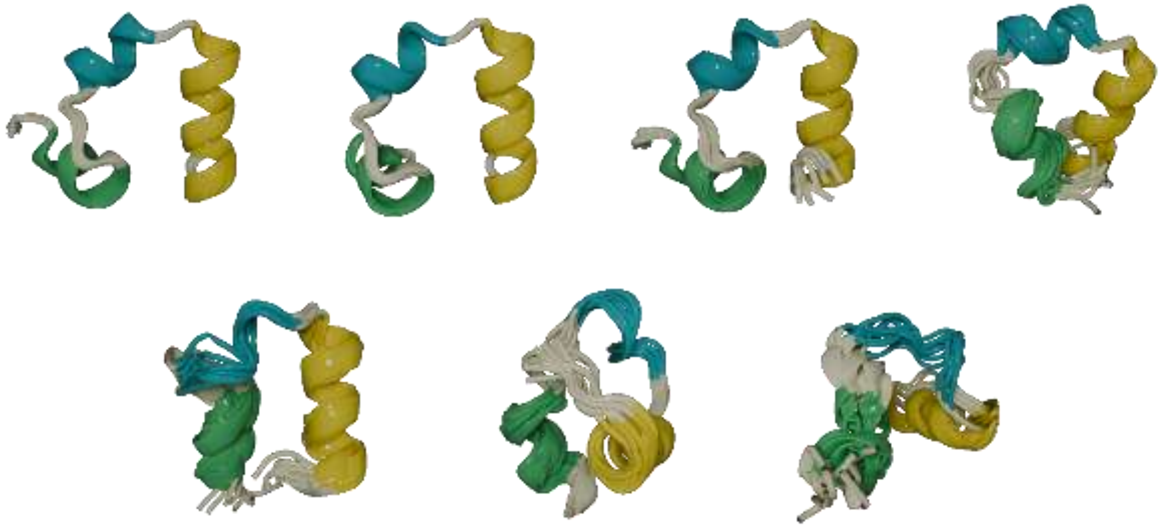

**Figure S1.** Overlaps of best representative structures from the seven clusters of HP35 after performing DIVINE using outlier_pair as anchors with refinement and weighted_MSD as cluster selection strategy. Helices 1, 2, and 3 are colored green, cyan, and yellow, respectively.

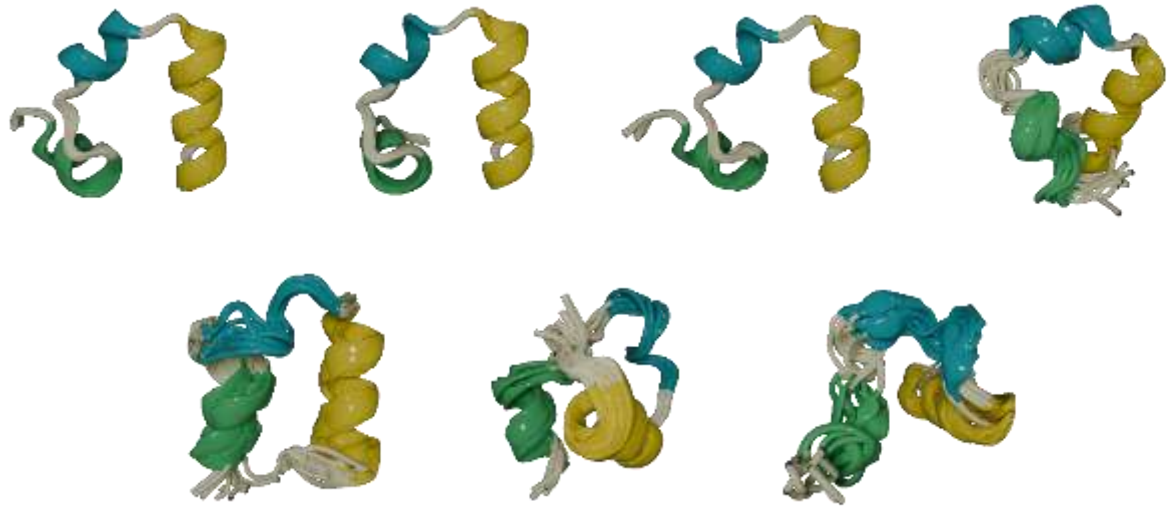

**Figure S2**. Overlaps of best representative structures from the seven clusters of HP35 after performing DIVINE using splinter_split as anchors with refinement and weighted_MSD as cluster selection strategy. Helices 1, 2, and 3 are colored green, cyan, and yellow, respectively.

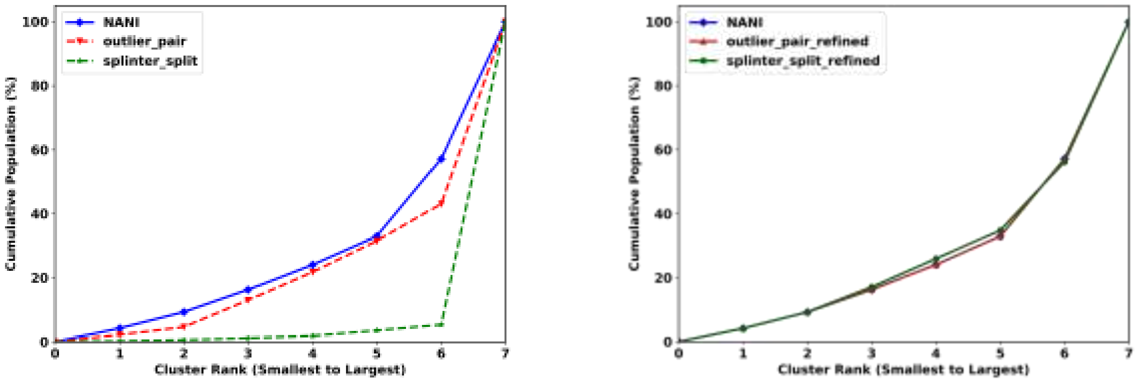


A B

**Figure S3.** Cumulative population distribution of clusters at $k = 7$ from DIVINE runs. (A) Unrefined anchor strategies (outlier_pair and splinter_split) against NANI. (B) Refined anchor strategies (outlier_pair_refined and splinter_split_refined) against NANI.

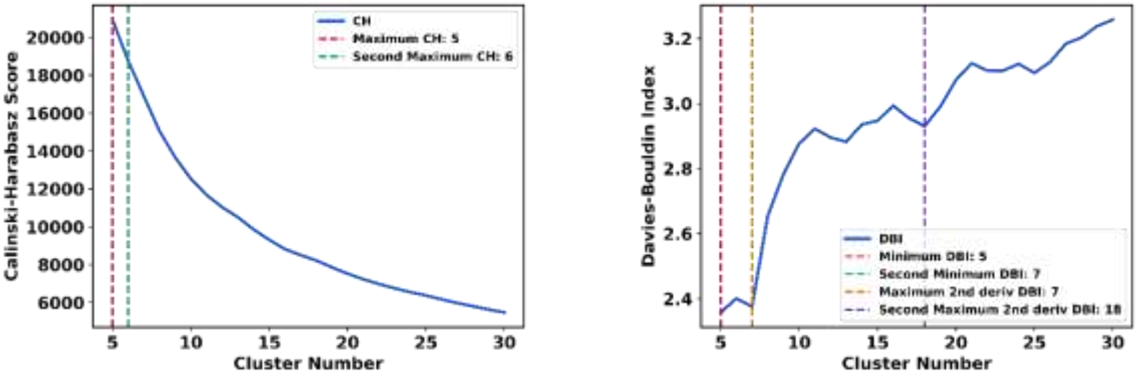


A B

**Figure S4.** Global and local analysis of clustering validity metrics for HP35 using DIVINE (NANI anchors, weighted_MSD). (A) Calinski-Harabasz and (B) Davies-Bouldin indices. Vertical dashed lines indicate the identified optima and significant inflection points (derived from second derivatives) within the analyzed range ($k \geq 5$).

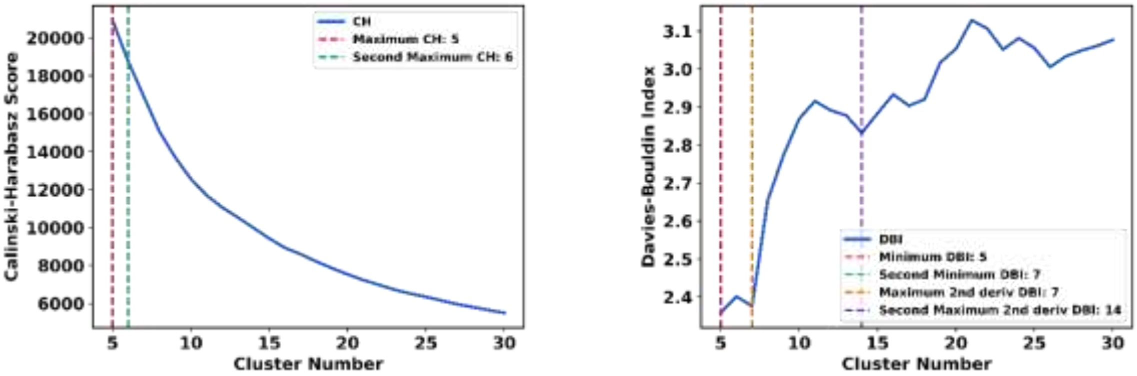


A B

**Figure S5.** Global and local analysis of clustering validity metrics for HP35 using DIVINE (outlier_pair_refined anchors, weighted_MSD). (A) Calinski-Harabasz and (B) Davies-Bouldin indices. Vertical dashed lines indicate the identified optima and significant inflection points (derived from second derivatives) within the analyzed range ($k \geq 5$).

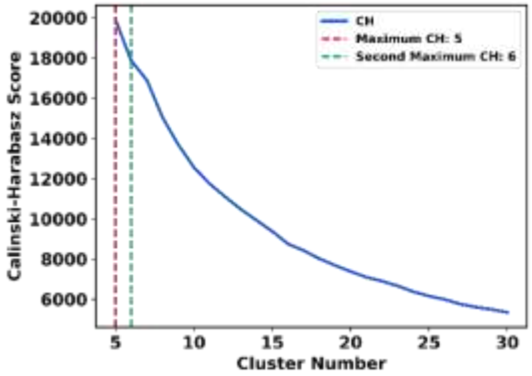


A

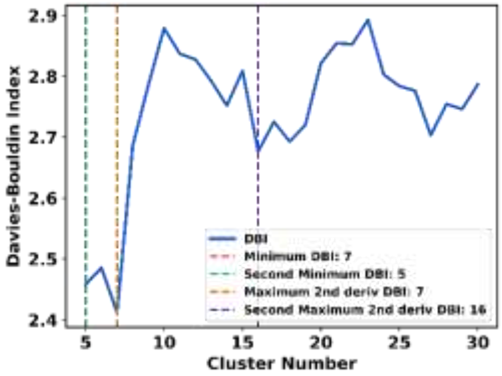


B

**Figure S6.** Global and local analysis of clustering validity metrics for HP35 using DIVINE (splinter_split_refined anchors, weighted_MSD). (A) Calinski-Harabasz and (B) Davies-Bouldin indices. Vertical dashed lines indicate the identified optima and significant inflection points (derived from second derivatives) within the analyzed range ($k \geq 5$).

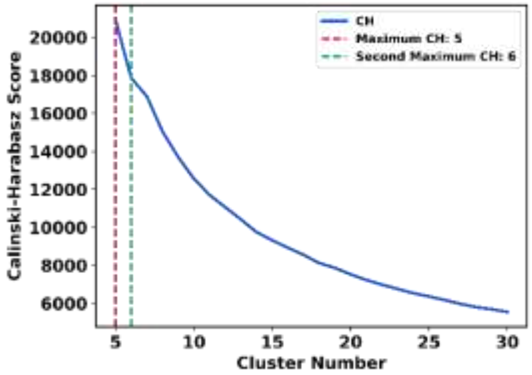


A1

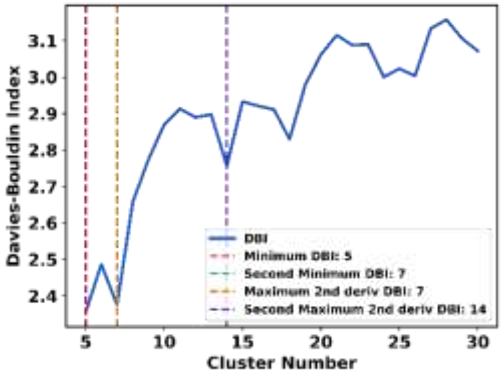


B1

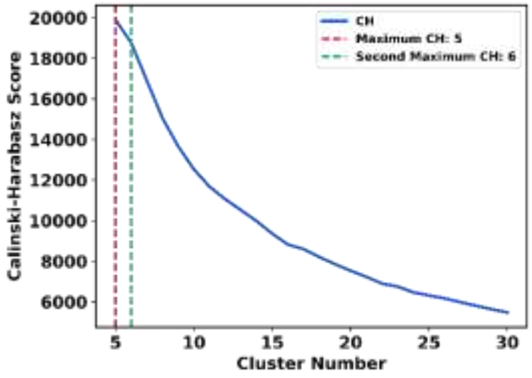


A2

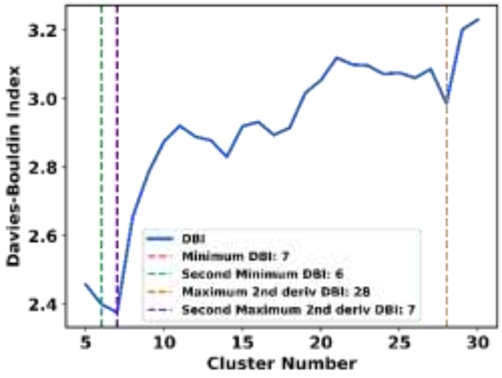


B2

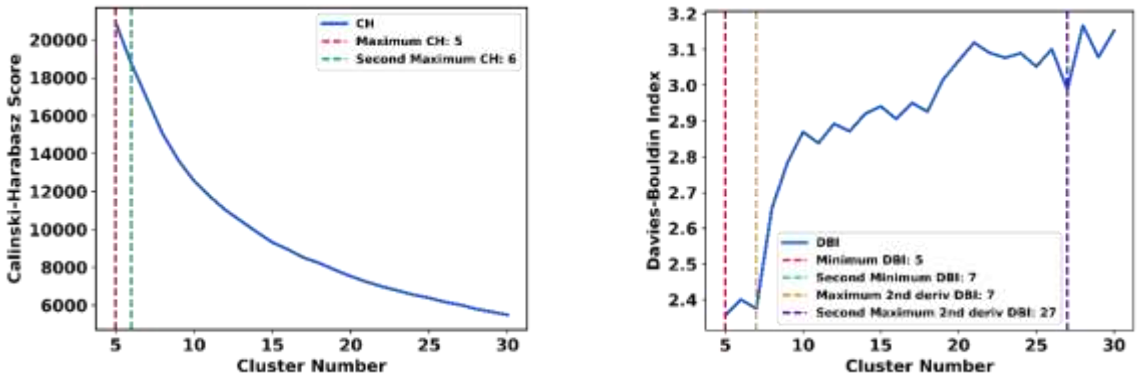


A3 B3

**Figure S7.** Global and local analysis of clustering validity metrics for three independent replicates of Bisecting $K$-means (BKM) (Biggest Inertia, k-means++ initialization) on HP35. (A1–A3) Calinski-Harabasz Index for Replicates 1, 2, and 3. (B1–B3) Davies-Bouldin Index for Replicates 1, 2, and 3. Vertical dashed lines indicate the identified optima and significant inflection points (derived from second derivatives) within the analyzed range ($k \geq 5$).

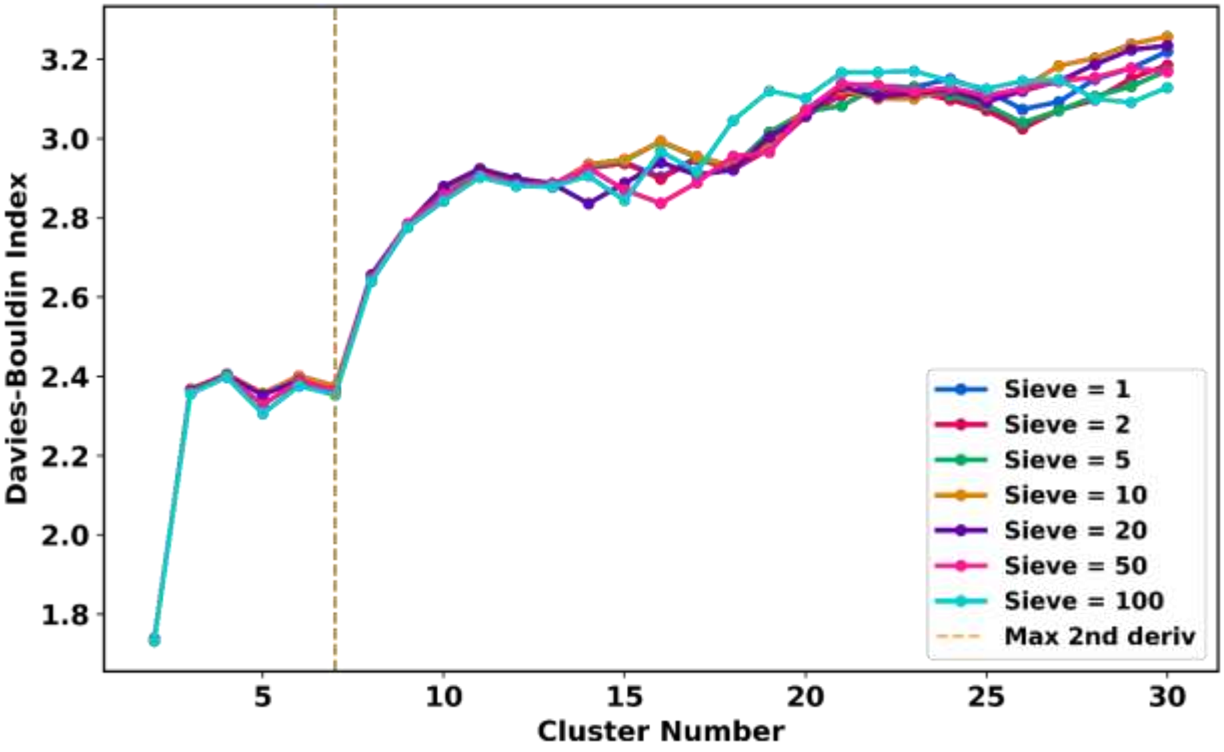


**Figure S8.** Robustness of clustering metrics across varying sampling densities. Davies-Bouldin Index profiles for HP35 clustering using DIVINE (NANI anchors, weighted_MSD) with trajectory sieving intervals of 1, 2, 5, 10, 20, 50, and 100 (corresponding to total frame counts ranging from ~1.5 million to ~15,000). Despite the reduction in data density, the profile shape and the location of the critical inflection point at $k = 7$ remain consistent.
